# Senolytic Treatment Reduces Cell Senescence and Necroptosis in Sod1 Knockout Mice that is Associated with Reduced Inflammation and Hepatocellular Carcinoma

**DOI:** 10.1101/2022.05.15.491998

**Authors:** Nidheesh Thadathil, Ramasamy Selvarani, Sabira Mohammed, Evan H. Nicklas, Albert L. Tran, Maria Kamal, Wenyi Luo, Jacob L. Brown, Marcus M. Lawrence, Agnieszka K. Borowik, Benjamin F. Miller, Holly Van Remmen, Arlan Richardson, Sathyaseelan S. Deepa

## Abstract

The goal of this study was to test the role cellular senescence plays in the increase in inflammation, chronic liver disease, and hepatocellular carcinoma, which are seen in mice null for Cu/Zn-Superoxide dismutase (Sod1KO). To inhibit senescence, six-month-old wildtype (WT) and Sod1KO mice were given the senolytics, dasatinib and quercetin (D+Q) for seven months. D+Q treatment reduced the expression of p16 in the livers of Sod1KO mice to WT levels as well as the expression of several SASP (senescence associated secretory phenotype) factors (IL-6, IL-1β, CXCL-1, and GDF-15). D+Q treatment also reduced markers of inflammation in livers of the Sod1KO mice, e.g., cytokines, chemokines, macropthage levels, and Kupffer cell clusters. D+Q treatment had no effect on various markers of liver fibrosis in the Sod1KO mice but reduced the expression of genes involved in liver cancer (Myc, Tgfbr2, Socs3, and Cdkn2a) as well as dramatically reducing the incidence of hepatocellular carcinoma. Surprisingly, D+Q also reduced markers of necroptosis (phosphorylated and oligomerized MLKL) in the Sod1KO mice to WT levels. We also found that inhibiting necroptosis in the Sod1KO mice with necrostatin-1s reduced the markers of cellular senescence (p16, p21, and p53). The data from our study suggest that an interaction occurs between cellular senescence and necroptosis in the liver of Sod1KO mice. We propose that these two cell fates interact through a positive feedback loop resulting in a cycle amplifying both cellular senescence and necroptosis leading to inflammaging and age-associated pathology in the Sod1KO mice.

## 1 INTRODUCTION

Cu/Zn-Superoxide dismutase is the major superoxide dismutase isozyme catalyzing the conversion of superoxide anions to hydrogen peroxide. It is found in all cells and is localized in the cytosol and the intermembrane space of the mitochondria (Okado-Matsumoto and Fridovich, 2001). In 1996, Reaume et al. generated mice null for Cu/Zn-Superoxide dismutase (Sod1KO) and reported that the Sod1KO mice appear normal at birth. However, the Sod1KO mice have very high levels of oxidative stress as measured by oxidative damage in various tissues and plasma (Muller et al., 2006). In addition, the lifespan of the Sod1KO mice was ∼25% shorter than wild type (WT) mice, with the median lifespan reduced from ∼30 months for WT mice to ∼22 months for Sod1KO mice (Zhang et al., 2013; Elchuri et al., 2005). Early studies showed that the Sod1KO mice exhibited various accelerated aging phenotypes, such as hearing loss (Keithley et al., 2005), cataracts (Olofsson et al. 2007), skin thinning and delayed wound healing (Iuchi et al. 2010), and sarcopenia (Muller et al. 2006). Our group showed that the Sod1KO mice also exhibited an accelerated loss of physical performance compared to age-matched WT mice as measured by voluntary running wheel activity, rota-rod performance, endurance exercise capacity, and grip strength (Deepa et al., 2017). More recently, we found that the overall severity of pathological lesions of Sod1KO mice was dramatically increased in adult Sod1KO mice compared to WT mice as measured by the geropathology grading score (Snider et al., 2018) and that the Sod1KO mice showed an accelerated decrease in cognition compared to WT mice (Logan et al., 2019). These data strongly suggest that the Sod1KO mice are a model of accelerated aging. The Sod1KO mice also develop chronic liver disease, such as fatty liver and liver fibrosis (Uchiyama et al., 2006; Sakiyama et al., 2016) and hepatocellular carcinoma, which is a major end-of-life pathology for the Sod1KO mice (Zhang et al., 2013; Elchuri et al., 2005).

We found that Sod1KO mice show a dramatic increase in circulating pro-inflammatory factors as well as increased markers of inflammation in various tissues (Mohammed et al., 2021a; Zhang et al., 2017). Because chronic, sterile inflammation (inflammaging) is believed to play an important role in aging and many age-related diseases, such as cancer, we proposed that increased inflammation in the Sod1KO mice was an important factor in the accelerated aging phenotype (Zhang et al., 2017). One of the pathways believed to play a role in inflammaging is cellular senescence (Wiley & Campisi, 2021; Olivieri et al., 2018). We found that markers of cellular senescence (p16, p21, and SA-βGal) were increased in kidney tissue from Sod1KO mice and reduced by dietary restriction (Zhang et al., 2017), which increased the lifespan of Sod1KO mice (Zhang et al., 2013). Importantly, the changes in p16 and p21 were associated with changes in the expression of proinflammatory cytokines (e.g., IL-6, and IL-1β) in kidney and circulating cytokines (Zhang et al., 2017). Based on these data we proposed that cellular senescence may be an important contributing factor to the accelerated aging phenotype observed in Sod1KO mice.

Recently, our attention turned to the potential role of necroptosis in inflammaging because it is a form of cell death that is highly proinflammatory due to the release of cell debris and self-molecules, damage-associated molecular patterns (DAMPs) (Pasparakis & Vandenabeele, 2015). We found that necroptosis was dramatically increased in the livers of Sod1KO mice, was associated with the increased inflammation and chronic liver disease observed, and treating Sod1KO mice with necrostatin-1s (Nec-1s), an inhibitor of necroptosis (Takahashi et al., 2012), reduced necroptosis and markers of inflammation in the liver (Mohammed et al., 2021a).

To gain a better understanding of the role cellular senescence plays in increased inflammation in Sod1KO mice, we treated Sod1KO mice with senolytics (dasatinib and quercetin, D+Q) to eliminate senescence cells (Zhu et al., 2015). We found that D+Q reduced inflammation and hepatocellular carcinoma in livers of the Sod1KO mice; however, we also found that D+Q also reduced markers of necroptosis in the livers of Sod1KO mice.

## 2 RESULTS

### 2.1 D+Q treatment reduces markers of cellular senescence in livers of Sod1KO mice

Female Sod1KO and WT mice were given dasatinib and quercetin (D+Q) for seven months starting at six months of age using the protocol developed by Kirkland’s group that was shown to eliminate sensescence cells *in vivo* (Zhu et al., 2015) and extend the healthspan and lifespan of 20-month-old WT mice (Xu et al., 2018). In this study, we focused on liver because cellular senescence and inflammation are increased in the livers of Sod1KO mice (Mohammed et al., 2021a) and because the Sod1KO mice develop fibrosis and hepatocellular carcinoma, which could arise from the increased cellular senescence and inflammation. Figure 1A shows that p16 and p21 transcripts are increased significantly in the livers of the Sod1KO mice as we have previously reported in male Sod1KO mice (Mohammed et al., 2021a). D+Q treatment significantly reduced the level of p16 transcripts to that observed in the WT mice; however, the the expression of p21 was not significantly reduced by D+Q treatment. We also measured the expression of twelve of the senescent associated secretory phenotype (SASP) factors commonly expressed by senescent cells (Acosta et al., 2008; Coppe et al., 2008). D+Q significantly reduced the expression of IL-6, IL-1β, CXCL-1, and GDF-15 in the livers of Sod1KO mice (Figure 1B, C).

**FIGURE 1.**
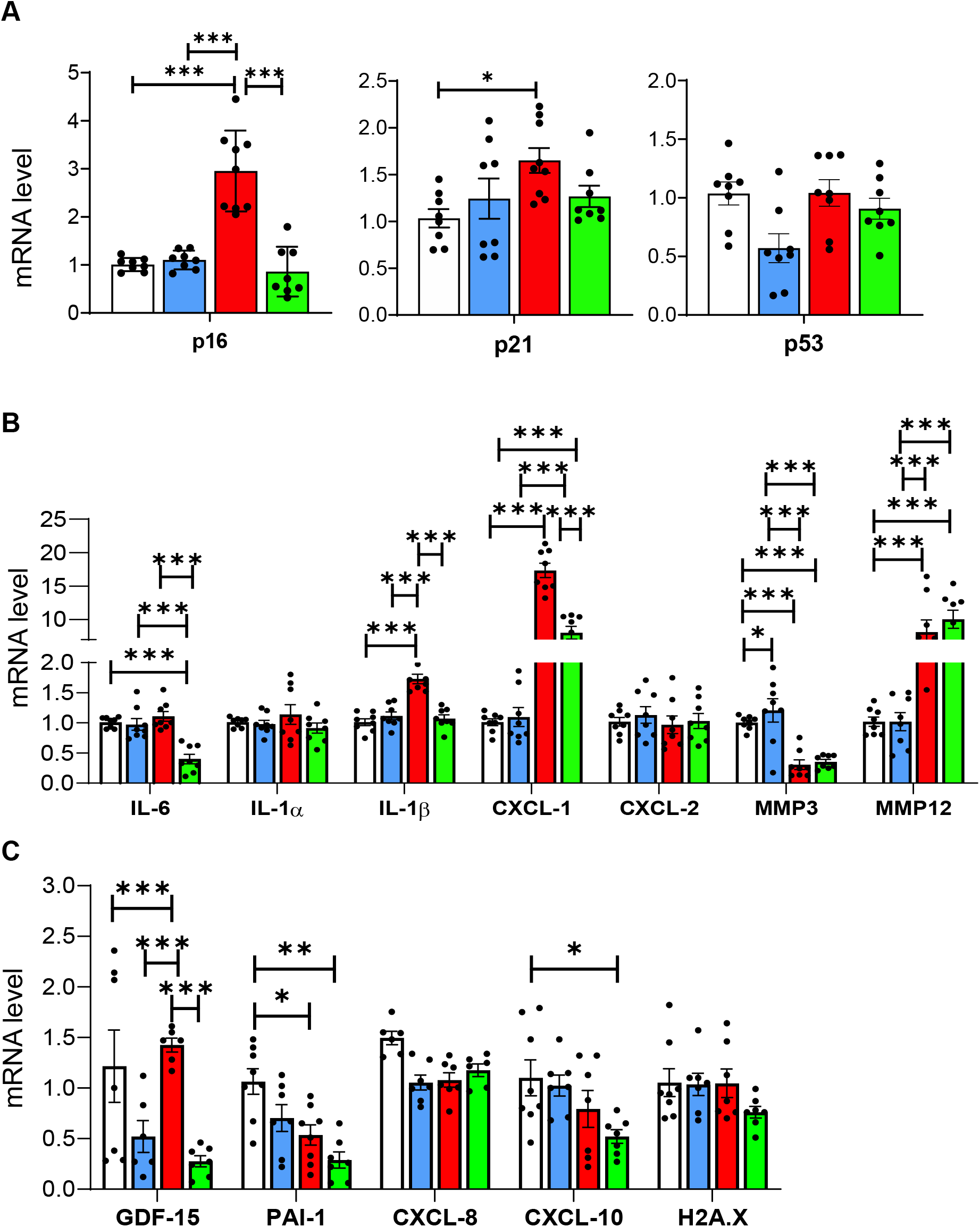
D+Q treatment significantly reduces markers of senescence and SASP-factors in the livers of Sod1KO mice. Panel **A** shows transcript levels of p16^Ink4a^ (Cdkn2a), p21^Cip1^ (Cdkn1a), and p53. Panels **B** and **C** shows the transcript levels of SASP-factors. The data in the graphs are generated using liver tissue from WT mice treated with vehicle (white bars), WT mice treated with D+Q (blue bars), Sod1KO treated with vehicle (red bars), and Sod1KO mice treated with D+Q (green bars). Data were obtained from 6 to 8 mice per group and are expressed as the mean ± SEM. (ANOVA, *** P ≤ 0.0005, ** P ≤ 0.005, * P≤ 0.05)

### 2.2 D+Q treatment reduces markers of inflammation in the livers of Sod1KO mice

We next studied the effect of D+Q treatment on markers of inflammation in the liver because cellular senescence has been associated with increased inflammation (Wiley & Campisi, 2021; Olivieri et al., 2018) and because Sod1KO mice show an increase in various markers of inflammation in liver (Mohammed et al., 2021a) and kidney (Zhang et al., 2017). Figure 2A shows the heat map for 84 proinflammatory cytokines and chemokines expressed in the livers of WT and Sod1KO mice treated with vehicle or D+Q. The data generated from the mouse cytokines and chemokines array are included in Table 1S in the supplement. It is evident that liver tissue from Sod1KO mice showed a dramatic increase in a large number of cytokines and chemokines compared to WT mice and that D+Q reduced the expression of most of these cytokines and chemokines. Figure 2B shows the quantification of the expression of ten of the chemokines that showed a statistically significant increase in expression in the livers of the Sod1KO mice. The expression of nine of the ten chemokines was significantly reduced (from 9 to 70%) by D+Q treatment. Figure 2C shows the quantification of the transcript levels of the seven cytokines that were significantly increased in the livers of the Sod1KO mice. D+Q treatment reduced the expression of all seven cytokines, and six of the cytokines were reduced to levels not significantly different from WT mice. Because macrophages are a major source of cytokines (Ju & Tacke, 2016) and because we have shown that markers of macrophages are increased in the livers of Sod1KO mice (Mohammed et al., 2021a), we measured macrophage levels in the livers of WT and Sod1KO mice treated with vehicle or D+Q. Figure 2D shows the transcript levels of F4/80, a marker of macrophage levels. F4/80 transcript levels were increased in the Sod1KO mice and reduced by D+Q. The accumulation of clusters of Kupffer cells is also a measure of proinflammatory status of liver (Woltman et al., 2014). We histologically compared liver tissue from the four groups of mice for the presence of Kupffer cell clusters (Figure 2E). Few if any Kupffer cell clusters were observed in liver tissue from the WT mice; however, we observed Kupffer cell clusters in the livers of the Sod1KO mice that was reduced in the Sod1KO mice treated with D+Q.

**FIGURE 2.**
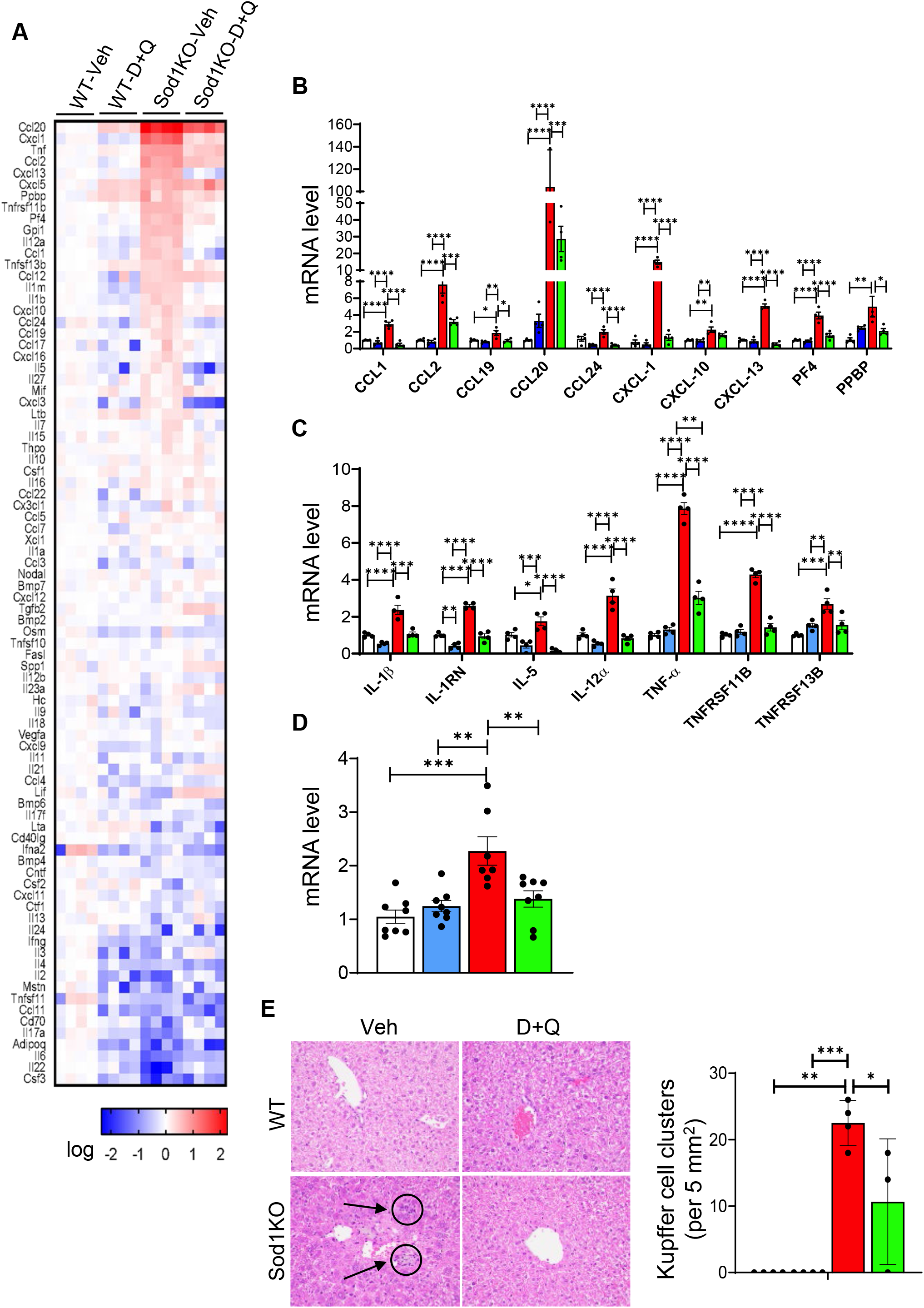
D+Q treatment significantly reduces markers of inflammation in the livers of Sod1KO mice. Panel **A** shows the heat map for the transcript levels of 84 cytokines and chemokines measured by the RT² Profiler™ PCR Array for liver tissue from WT and Sod1KO mice treated with vehicle or D+Q. Transcript levels greater than the mean are shaded in red and those lower than the mean are shaded in blue. Panel **B** shows the expression of the ten chemokines, and Panel **C** shows the expression of the seven cytokines from the Profiler™ PCR Arrays that are significantly increased in the livers of the Sod1KO mice. The data were obtained from 4 to 6 mice per group and are expressed as mean ± SEM. Panel **D** shows the transcript levels of macrophage marker F4/80 in the livers of WT and Sod1KO mice treated with vehicle and/or D+Q. Data were obtained from 7 to 8 mice per group and are expressed as mean ± SEM. Panel **E** shows the detection of Kupffer cells cluster by H & E staining (left), and quantification of Kupffer cell clusters in the livers WT and *Sod1*KO mice treated with vehicle or D+Q (right). Data were obtained from 3 to 5 mice per group and are expressed as the mean ± SEM. The data in the graphs are shown as follows: white bars for WT mice treated with vehicle, blue bars for WT mice treated with D+Q, red bars for Sod1KO treated with vehicle, and green bars for Sod1KO mice treated with D+Q. (ANOVA, **** P < 0.0001, *** P ≤ 0.0005, ** P ≤ 0.005, * P≤ 0.05).

### 2.3 D+Q treatment has no impact on fibrosis in the livers of Sod1KO mice

We next determined if D+Q treatment had an impact on chronic liver disease, which is increased in the Sod1KO mice (Uchiyama et al., 2006; Sakiyama et al., 2016; Mohammed et al., 2021a). The data in Figure 3B shows that circulating alanine aminotransferase (ALT) levels, which is a measure of liver damage, are increased in the Sod1KO mice as previoulsy observed (Mohammed et al., 2021a). However, ALT levels were not significantly altered by D+Q treatment. We next measured various markers of fibrosis in the four groups of mice. The activation and proliferation of stellate cells play a major role in the pathogenesis of hepatic fibrosis (Guido et al., 1996), and activated stellate cells up regulate the expression of desmin (Kim et al., 2012). Figure 3A shows that desmin levels are increased over 5-fold in liver tissue from Sod1KO mice compared to WT mice; however, D+Q treatment did not significantly reduce desmin levels in the Sod1KO mice. We also measured other markers of fibrosis in liver tissue from the four groups of mice: transcript levels of collagen 1α1 (Col1α1), hydroxyproline levels, and histopathological changes. Col1α1 expression (Figure 3C) and hydroxyproline levels (Figure 3D) were significantly increased in the livers of the Sod1KO mice, and D+Q treatment did not reduce these two markers of liver fibrosis. Histopathological evidence of liver fibrosis was assessed by the appearance of perisinusoidal/pericellular (chicken wire) fibrosis using Picrosirius Red staining (Figure 3E. This marker of fibrosis was incresed in the livers of Sod1KO mice, and D+Q treatment had no significant effect on this pathological lesion. Thus, our data show that D+Q treatment had no impact on the increase in fibrosis observed in the Sod1KO mice.

**FIGURE 3.**
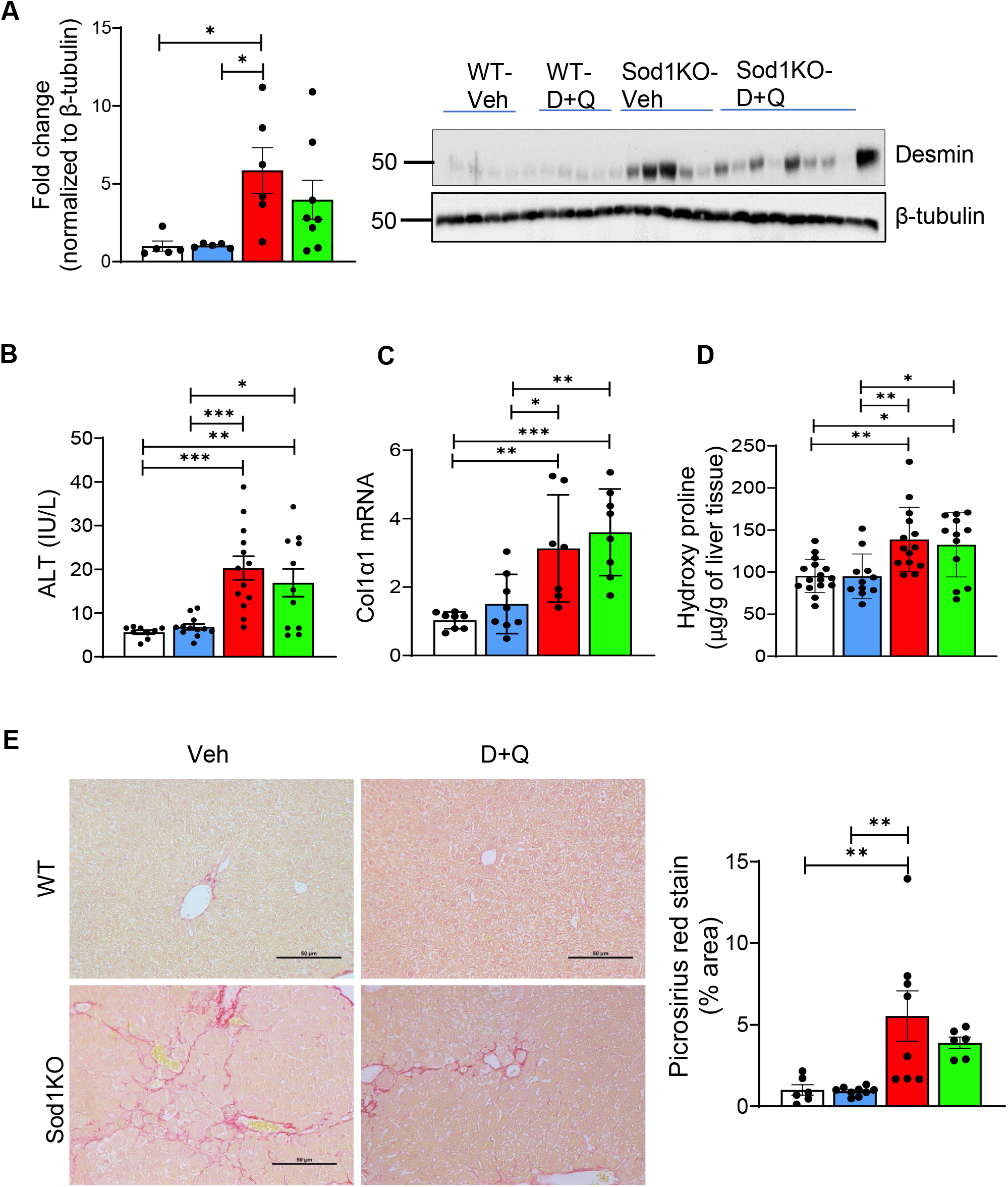
D+Q treatment does not reduce fibrosis in the livers of Sod1KO mice. Panel **A** shows the quantification of desmin by western blots normalized to β-tubulin. Panel **B** shows the serum levels of alanine aminotransferase (ALT) expressed as IU/L. Panel **C** shows transcript levels of Col1α1 normalized to β-microglobulin and expressed as fold change. Panel **D** shows hydroxyproline levels expressed as µg of hydroxyproline/g of liver tissue. Panel **E** shows sections from Picrosirius Red staining (scale bar: 50µm) (left) and the quantification of fibrotic area (right) for livers from WT and Sod1KO mice treated with vehicle or D+Q. The data in the graphs are shown as follows: white bars for WT mice treated with vehicle, blue bars for WT mice treated with D+Q, red bars for Sod1KO treated with vehicle, and green bars for Sod1KO mice treated with D+Q. The data were obtained from 6 to 12 mice per group and are expressed as the mean ±SEM. (ANOVA, *** P ≤ 0.0005, ** P ≤ 0.005, * P≤ 0.05).

### 2.4 D+Q treatment reduces hepatocellular carcinoma in the livers of Sod1KO mice

Because hepatocellular carcinoma is a major pathology observed in Sod1KO mice (Elchuri et al., 2005; Zhang et al., 2013), we studied the impact of D+Q on the development of cancer in the livers of the Sod1KO mice. The increase in liver weight, which is characteristic of the Sod1KO mice and associated with the development of liver tumors, was reduced by D+Q treatment (Figure 4A) as was the increase in circulating levels of α-fetoprotein (AFP), a liver tumor marker (Figure 4B). Figure 4C shows the heat map for 84 genes involved in the progression of hepatocellular carcinoma as well as other forms of hepatocarcinogenesis. The data generated from the liver cancer array are included in Table 2S in the supplement. These genes are involved in commonly altered signal transduction pathways in cancer as well as genes involved in other dysregulated biological pathways such as epithelial to mesenchymal transition, cell cycle, apoptosis, and inflammation. In addition to liver tissue from the WT mice and and non-tumor tissue from Sod1KO mice, we also studied tumor tissue from the Sod1KO mice. The expression of twenty genes were significantly up-regulated in tumor tissue from Sod1KO mice, and the expression of many of these genes was increased in non-tumor tissue from the Sod1KO mice. Figure 4D shows the expression of four genes that have been shown to play a role in liver cancer that were significantly increased in non-tumor and tumor tissue from Sod1KO mice: Myc, Tgfbr2, Socs3, and Cdkn2a. D+Q reduced the expression of these genes to the basal levels observed in the WT mice. We carefully checked the liver tissue from each of the mice used in the study for the presence of tumors, and as shown in Figure 1S in the supplement, these tumors expressed markers specific for hepatocellular carcinoma, e.g., glypican 3 and α-fetoprotein (Zhao et al., 2013). Figure 4E shows the incidence of tumors in the WT and Sod1KO mice. As expected, we observed no tumors in any of the WT mice. However, 64% (7 out of 11) of the Sod1KO mice showed the presence of liver tumors. D+Q dramatically reduced the incidence of liver tumors in the Sod1KO mice to only 1 out of the 13 Sod1KO mice treated with D+Q.

**FIGURE 4.**
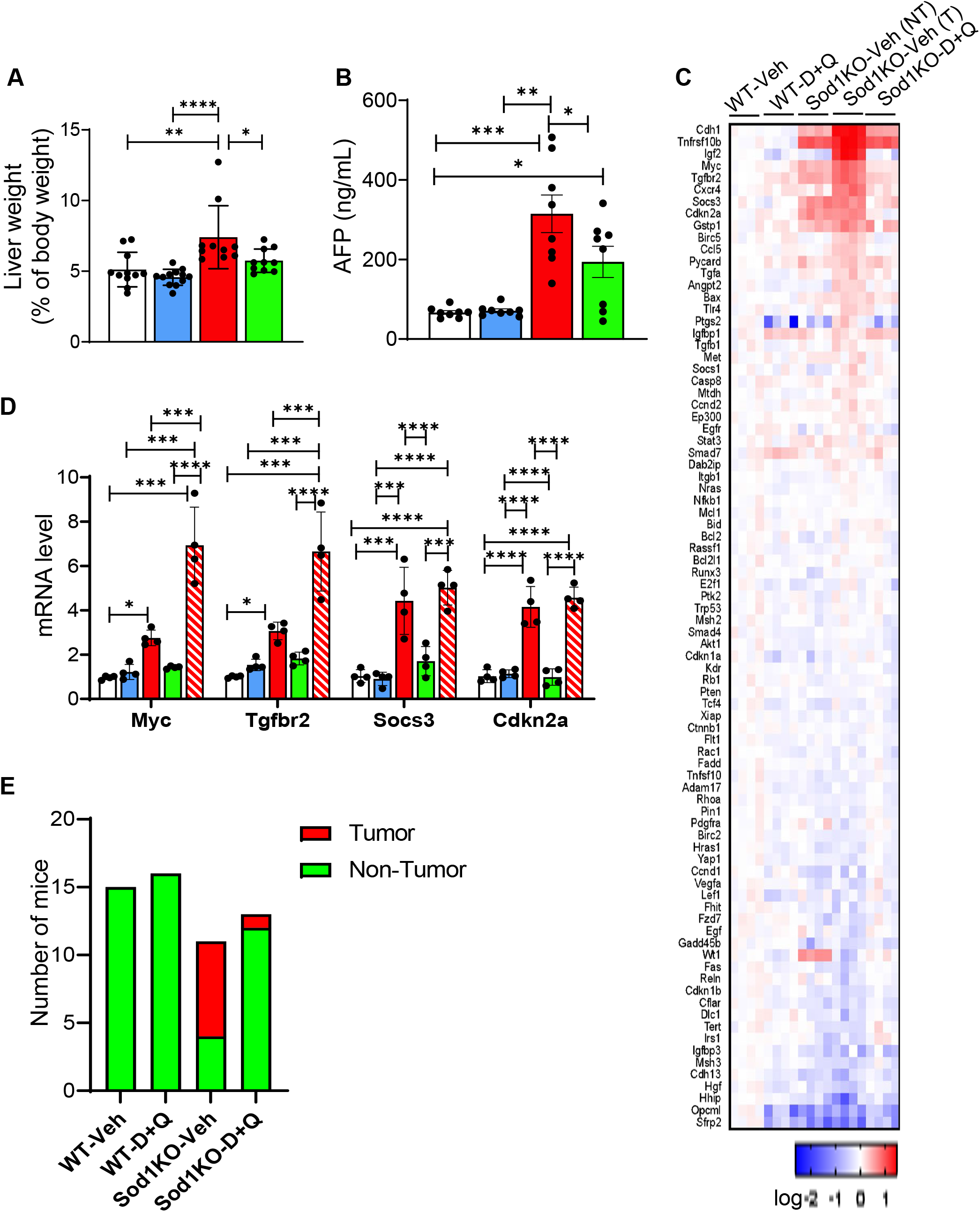
Liver cancer is reduced in Sod1KO mice treated with D+Q. Panel **A** shows the liver weight normalized to body weight. Panel **B** shows α-fetoprotein (AFP) levels in the serum expressed as ng/mL. Panel **C** shows the heat map for the RT² Profiler™ PCR Arrays of 84 genes related to liver cancer for normal liver tissue from WT and Sod1KO mice treated with vehicle or D+Q as well as tumor tissue from Sod1KO mice treated with vehicle. Transcript levels greater than the mean are shaded in red and those lower than the mean are shaded in blue. Panel **D** shows the quantification of the fold change in mRNA levels for four genes (Myc, Tgfbr2, Socs3 and Cdkn2a) that were significantly increased in normal tissue from Sod1KO mice treated with vehicle. The data were obtained from 4 to 6 mice per group and are expressed as mean ± SEM. The data in the graphs in Panels **A**, **B** and **D** are shown as follows: non-tumor tissue from WT mice treated with vehicle (white bars), WT mice treated with D+Q (blue bars), Sod1KO treated with vehicle (red bars), and Sod1KO mice treated with D+Q (green bars) and tumor tissue from Sod1KO mice treated with vehicle (red bars with strips). (ANOVA, **** P < 0.0001, *** P ≤ 0.0005, ** P ≤ 0.005, * P≤ 0.05). Panel **E** graphically shows the number of WT and Sod1KO mice treated with vehicle or D+Q that showed the presence (red) or absence of hepatocellular carcinoma tumors (green). None of the WT mice treated with vehicle (n=15) or D+Q (n=11) showed the presence of tumors while seven of Sod1KO mice treated with vehicle (n=11) and one of the Sod1KO mice treated with D+Q (n=13) showed the presence of tumors.

### 2.5 D+Q treatment reduces markers of necroptosis in the livers of Sod1KO mice

We also studied the impact of D+Q treatment on other cell-fates because D+Q has been shown to alter apoptosis (Kirkland & Tchkonia, 2020) and because we have shown that necroptosis is increased in liver tissue from Sod1KO mice (Mohammed et al., 2021a). As shown in Figure 5A, we observed no significant change in cleaved caspase 3 activity, a marker of apoptosis, in the livers from the WT or Sod1KO mice. Next, we measured the level of necroptosis in liver tissue from the WT and Sod1KO mice by measuring the phosphorylation and oligomerization of MLKL, which is the terminal step in necroptosis resulting in the disruption of the cell membrane and release of DAPMs (Chen et al., 2014; Huang et al., 2017). A dramatic increase in P-MLKL levels (Figure 5B) and MLKL oligomerization (Figure 5C) was observed in the livers of the Sod1KO mice, which is consistent with our previously published data (Mohammed et al., 2021a). To our surprise, we observed that D+Q treatment reduced these markers of necroptosis in the Sod1KO mice to the levels observed in WT mice. These data suggest that either D+Q inhibits necroptosis as well as cell senescence or that inhibiting cell senescence leads to reduced necroptosis, i.e., there is an interaction between cell senescence and necroptosis where inhibiting one cell-fate impacts the other. To test the second possibility, we studied whether inhibiting necroptosis in the Sod1KO mice had an effect on cellular senescence. In these experiments, we used liver tissue from male Sod1KO mice used in a previous study, in which we showed that 25 days of necrostatin-1s (Nec-1s) treatment reduced necroptosis (Mohammed et al., 2021a). Figure 5D shows the expression of p16, p21, and p53 in liver tissue from these Sod1KO mice. Nec-1s treatment reduced p16, p21, and p53 transcript levels, demonstrating that inhibiting necroptosis in Sod1KO mice is also associated with reduced senescence.

**FIGURE 5.**
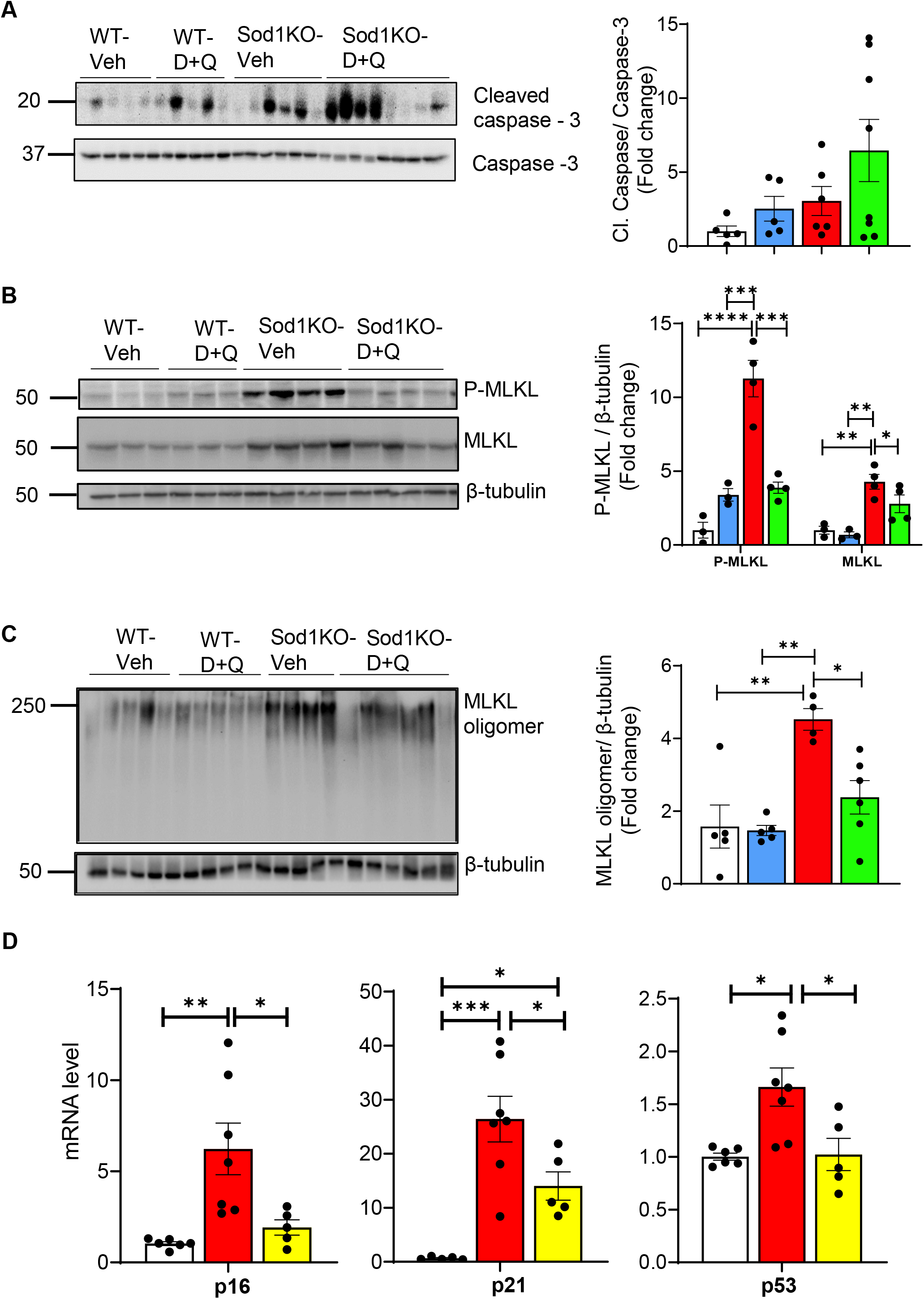
The impact of D+Q treatment on apoptosis and necroptosis and Nec-1s treatment on cellular senescence in the livers of Sod1KO mice. Panel **A** shows western blots of cleaved caspase-3 and caspase-3 (left) and the quantification of cleaved caspase-3 normalized to caspase-3 (right). Panel **B** shows western blots of P-MLKL, MLKL, and β-tubulin (left) and the quantification of P-MLKL and MLKL normalized to β-tubulin (right). Panel **C** shows western blots of MLKL oligomers and β-tubulin (left) and the quantification of MLKL oligomers normalized to β-tubulin (right). The data in the graphs in Panels **A**, **B**, and **D** are shown as follows: liver tissue from for WT mice treated with vehicle (white bars), WT mice treated with D+Q (blue bars), Sod1KO treated with vehicle (red bars), and Sod1KO mice treated with D+Q (green bars). Panel **D** show the effect of Nec-1S treatment on markers of cellular senescence that was obtained from the livers of male WT and Sod1KO mice studied by Mohammed et al. (2021a). The data in the graphs are shown as follows. WT treated with vehicle (white bars), Sod1KO treated with vehicle (red bars), and Sod1KO mice treated with Nec-1s (yellow bars). The data in all the graphs were obtained from 4 to 6 mice per group and are expressed as the mean ± SEM. (ANOVA, **** P < 0.0001, *** P ≤ 0.0005, ** P ≤ 0.005, * P≤ 0.05).

## 3 DISCUSSION

Chronic, low-grade inflammation that occurs with age (inflammaging) has been identified as one of the ‘seven pillars of aging’ (Kennedy et al., 2014). Inflammaging has been observed in all mammalian species studied including rodents (Brubaker et al., 2011), rhesus monkeys (Didier et al., 2012), and humans (Franceschi & Campisi, 2014). Because inflammation is strongly associated with a variety of diseases, such as cardiovascular disease, cancer, type 2 diabetes, frailty, and neurodegenerative diseases, it has been argued that inflammaging is an important factor in the etiology of these diseases (Franceschi & Campisi, 2014; Ferrucci & Fabbri, 2018). In addition, studies with mice show that interventions that increase lifespan, such as dietary restriction (Spaulding et al., 1997), dwarfism (Masternak et al., 2012), and rapamycin treatment (Richardson et al., 2015) reduce inflammation. On the other hand, Sod1KO mice, which show accelerated aging, exhibit an increase in markers of inflammation (Zhang et al., 2016). At the present time, several pathways have been proposed to play a role in inflammaging, and cellular senescence is one of these pathways (Wiley & Campisi, 2021; Olivieri et al., 2018). Cytokines and chemokines produced and secreted by senescent cells as part of the SASP have the potential to activate the immune system resulting in the generation of an inflammatory response. We previously found that markers of cellular senescence were higher in the Sod1KO mice and were reduced by dietary restriction, which was associated with an increase or decrease, respectively, in markers of inflammation (Zhang et al., 2016). These data led us to propose that cellular senescence played a role in increased inflammation and the accelerated aging phenotype observed in the Sod1KO mice.

The goal of this study was to test whether cellular senescence plays a role in the increase in inflammation and the progression of fibrosis and hepatocellular carcinoma observed in Sod1KO mice by treating Sod1KO mice with senolytics to eliminate senescent cells. Using a combination of dasatinib (a tyrosine kinase inhibitor) and quercetin (a flavonoid), we found that seven months of D+Q treatment reduced the expression of p16 in the livers of Sod1KO mice to WT levels as well as reducing the transcript levels of several SASP-factors. Importantly, we observed that D+Q treatment reduced markers of inflammation in liver supporting our proposal that cellular senescence was at least partially responsible for the increased hepatic inflammation observed in the Sod1KO mice. Because inflammation has been shown to play a role in liver fiborsis (Duval F et al., 2015), we studied the effect of D+Q on fibrosis, which is increased in the livers of the Sod1KO mice (Uchiyama et al., 2006; Sakiyama et al., 2016; Mohammed et al., 2021a). We were surprised to find that D+Q treatment had no effect on the various markers of fibrosis we assessed. One possible explanation for the lack of an effect of D+Q on fibrosis was because the Sod1KO mice were six months of age when D+Q was first administered. Sakiyama et al. (2016) has shown that increased collagen deposition in the livers of Sod1KO mice occurs as early as three months of age. Therefore, fibrosis is already developed at six months of age in the Sod1KO mice and a reduction in cellular senescence and inflammation at this age does not reverse fibrosis.

In contrast to fibrosis, we found that D+Q treatment reduced the expression of several genes involved in liver cancer as well as dramatically reducing the incidence of hepatocellular carcinoma in the Sod1KO mice. Chronic inflammation has been shown to play an important role in hepatocellular carcinoma. For example, higher baseline serum levels of inflammatory markers (CRP, IL-6, C-peptide) are associated with an increased risk of developing hepatocellular carcinoma in the general population (Aleksandrova et al., 2014), and the use of anti-inflammatory drugs is linked to lower risk and better survival in patients with hepatocellular carcinoma (Tao et al., 2018; Pang et al., 2017; Sahasrabuddhe et al, 2012). In addition, the enhanced production of proinflammatory cytokines TNF-α and IL-6 are key mediators of obesity-mediated hepatocellular carcinoma (Park et al., 2010; Yu et al., 2018; Villanueva & Luedde, 2016). Therefore, our data are consistent with the reduction in inflammation and cellular senescence being responsible for the reduction of hepatocellular carcinoma observed in the D+Q treated Sod1KO. However, it is also possible that the reduction in hepatocellular carcinoma is due to a direct effect of dasatinib on the growth of cancer cells. Dasatinib is second generation tyrosine kinase inhibitor that downregulates various tyrosine kinases containing BCR-ABL1 and kinases of SRC family, and it inhibits the growth of cancer cells (Zhang et al., 2020), including hepatocellular carcinoma cells (Liu et al., 2021; Chang et al., 2013).

Because we previously observed that both cellular senescence (Zhang et al., 2017) and necroptosis (Mohammed et al., 2021a) were increased in the Sod1KO mice, our goal in this study was to use D+Q treatment to differentiate the effect of cellular senescence and necroptosis on the increased inflammation observed in the Sod1KO mice. We were surprised to find that D+Q also reduced markers necroptosis in the Sod1KO mice. Thus, the decrease in inflammation observed in the D+Q treated Sod1KO mice could be due to cellular senescence, necroptosis, or both. While this result was surprising, it was not totally unexpected because we recently found that Nec-1s treatment, which inhibits necroptosis, reduced markers of both necroptosis and cell senescence in the livers of old mice (Mohammed et al., 2021b). In Sod1KO mice, we found in this study that Nec-1s treatment reduced markers of cellular senescence in the liver. Thus, the data from our studies with D+Q and Nec-1s suggest that cellular senescence and necroptosis could be interacting through a positive feedback loop resulting in a cycle amplifying both cell-fates leading to inflammaging and age-associated pathology. Senescent cells, which accumulate with age, could trigger necroptosis in surrounding cells by the paracrine/juxtracrine actions of the SASP-factors (cytokines, chemokines, extracellular proteases, growth factors, lipids, etc.) secreted from senescent cells. For example, TNF-α, a component of SASP (Acosta et al., 2013; Freund et al., 2010) is a potent inducer of necroptosis (Liu et al., 2014; Vanlangenakker et al., 2011). Because senescent cells are resistant to apoptosis and accumulate with age, they are a potential source of factors that could push cells to undergo necroptosis in old animals or in Sod1KO mice leading to the release of DAMPs, which are potent inducers of inflammation (Kataoka et al., 2014; Zhang et al., 2010). The elimination of senescent cells in Sod1KO mice by D+Q treatment could therefore reduce necroptosis. In turn, DAMPs arising from necroptotic cells could increase the burden of senescent cells by inducing cells to undergo senescence. The DAMPs, HMGB1 and IL-1 have been shown to induce cultured fibroblasts to undergo senescence (Acosta et al., 2013; Davalos et al., 2013). In addition, extracellular vesicles have been shown to induce cellular senescence (Borghesan et al., 2019), and extracellular vesicles are released during necroptosis (Yoon et al., 2017). Therefore, reducing necroptosis by Nec-1s could reduce the potential of DAMPs inducing cells to undergo senescence.

In summary, our study shows for the first time that the senolytics, D+Q not only reduce cellular senescence but also reduce necroptosis. We propose that the reduction in necroptosis by D+Q most likely arises from a reduction in the number of senescent cells that secrete SASP-factors such as TNF-α, which induce cells to undergo necroptosis. Therefore, cell senescence and necroptosis could play an important role in inflammaging resulting in the progression of many age-related diseases and aging. Whether the same or different cell type(s) in the liver undergo cell senenscence and necroptosis in Sod1KO mice is currently under investigation.

## 4 EXPERIMENTAL PROCEDURES

### 4.1 Animals

The Sod1KO and WT mice were generated and raised in animal facilities at the Oklahoma Medical Research Foundation (OMRF), and all procedures were approved by the IACUC at OMRF. The mice were group housed in ventilated cages at 20 ± 2 °C, on a 12-h/12-h dark/light cycle, and fed a commercial rodent chow diet *ad libitum*. Four groups of female mice were used in the study: WT mice treated with vehicle, WT mice treated with D+Q, Sod1KO mice treated with vehicle, and Sod1KO mice treated with D+Q. At six months of age, mice were given dasatinib (5mg/kg) and quercetin (50mg/kg) dissolved in vehicle (10% ethanol, 30% polyethylene glycol 400 and 60% Phosal 50 PG) or vehicle by oral gavage as described by Zhu et al. (2015) for three constitutive days every fifteen days over seven months. The final dose of D+Q was given seven days prior to sacrifice at thirteen months of age. Liver tissues was collected, immediately frozen in liquid nitrogen, and stored at -80°C until use in the experiments described below.

### 4.2 RNA isolation, cDNA synthesis, and quantitative real-time PCR

Real-time-PCR was performed using 20mg frozen liver tissues as described previously by Mohammed et al. (2021). Briefly, total RNA was extracted using RNeasy kit (Qiagen, Valencia, CA, USA) as per manufacturer’s instructions. First-strand cDNA was synthesized using a high-capacity cDNA reverse transcription kit (ThermoFisher Scientific, Waltham, MA), and quantitative RT-PCR was performed with Power SYBR Green PCR Master Mix (ThermoFischer Scientific Waltham, MA). The primers used for RT-PCR analysis are given in the Table 3S in the supplement. Calculations were performed by a comparative method (2^−ΔΔ^Ct) using β-microglobulin or β-actin as controls as described previously (Mohammed et al., 2021a).

Genes involved in liver cancer were analyzed by Mouse RT^2^ Profiler^TCM^ PCR Liver Cancer Array [Qiagen, Cat# PAMM-1332E-4 (330231)], which measures the expression of 84 genes involved in the progression of HCC, as well as other forms of hepatocarcinogenesis. Similarly, the RT² Profiler™ PCR Array Mouse Cytokines & Chemokines [Qiagen, Cat# PAMM-1502E-4 (330231)] was used to analyze the 84 cytokines and chemokines levels in the liver samples.

### 4.3 Western blotting

Western blots were performed as described previously (Mohammed et al., 2021a, b). Images were taken using ChemiDoc imaging system (Bio-Rad Laboratories, Hercules, CA) and quantified using ImageJ Software (U.S. National Institutes of Health, Bethesda, MD, USA). Primary antibodies against the following proteins were used: phospho(S345) MLKL from Abcam (Cambridge, MA); MLKL from Millipore Sigma (Burlington, MA); Cleaved Caspase-3, and Caspase from 3 Cell Signaling Technology (Danvers, MA); desmin from ThermoFisher Scientific (Waltham, MA); β-tubulin from Sigma-Aldrich (St. Louis, MO). HRP-linked anti-rabbit IgG, HRP-linked anti-mouse IgG and HRP-linked anti-rat IgG from Cell Signaling Technology (Danvers, MA) were used as secondary antibody.

### 4.4 MLKL oligomerization

MLKL oligomerization was measured as described by Miyata et al. (2021). Briefly, liver tissue was homogenized in HEPES buffer (pH 7.4), and protein in the homogenate quantified using Bradford method. Protein samples were prepared using 2x Laemmli buffer without any reducing agents to maintain the proteins under non-reducing conditions. Forty μg of protein was resolved under non-reducing conditions without SDS in running buffer and poly acrylamide gel. MLKL oligomers were detected on the gels as larger than 200 kDa and quantified using the antibody to MLKL as described above.

### 4.5 Histological examination of liver for clusters of Kupffer cells

Formalin fixed liver tissue was embedded in paraffin, and 4μm sections were generated using a microtome. Haematoxylin and eosin (H & E) staining was performed on the tissue samples using the standard procedure at the Stephenson Cancer Center Tissue Pathology Core. H & E stained sections were digitally scanned at 10X and 20X magnifications using Nikon Nikon Ti Eclipse microscope (Nikon, Melville, NY). Clusters of Kupffer cells in the liver tissue were identified by a pathologist, and the Kupffer cell clusters in tissue sections were measured in three random fields per sample and quantified using Image J software.

### 4.6 Immunohistochemistry analysis for hepatocellular carcinoma

Immunohistochemistry staining was performed as described by Thadathil et al. (2021) with modifications. Briefly, 4μm paraffin embedded liver sections were deparaffinized using graded series of xylene:ethanol, and the antigen retrieval was performed with Proteinase K treatment. Liver sections were then permeabilized, blocked, and incubated with primary antibodies against glypican-3 or α-fetoprotein (Thermofisher Scientific, Waltham, MA). After washing, sections were incubated with HRP conjugated anti-mouse or anti-rabbit secondary antibodies for 1 hour. The sections were then visualized using 3,3 diaminobenzidine followed by Mayer’s Hematoxylin (Sigma-Aldrich, St. Louis, MO) counter staining. These sections were washed, dehydrated with gradient ethyl alcohol, cleared in xylene, and mounted using DPX mounting media. The images were taken using a Nikon Ti Eclipse microscope (Nikon, Melville, NY) for 3 random fields per sample.

### 4.7 Alanine aminotransferase (ALT) assay

The activity of ALT in the serum was measured using the assay kit from Cayman Chemical Company (Ann Arbor, MI) following manufacturer’s instructions.

### 4.8 α-Fetoprotein (AFP) assay

Serum AFP levels were quantified using the mouse α-fetoprotein/AFP Quantikine ELISA Kit (R&D Systems, Minneapolis, MN) according to the manufacturer’s instructions.

### 4.9 Hydroxyproline assay

Hydroxyproline content was measured as described by Smith et al. (2016). Liver tissue (∼250mg) was pulverized using liquid nitrogen and was digested in 6 M hydrochloric acid overnight at 110°C. Ten ml of the digest was mixed with 150μl of isopropanol, 75μl of Solution A (1:4 mix of 7% Chloramine T (Sigma-Aldrich, St. Louis, MO), and acetate citrate buffer [containing 57g sodium acetate anhydrous, 33.4g citric acid monohydrate, 435ml 1M sodium hydroxide, 385ml isopropanol in 1L of buffer]. The mixture was vigorously mixed and incubated at room temperature for 10 minutes. After incubation, 1ml of Solution B [3:13 mix of Ehrlich’s reagent (3g p-dimethyl amino benzalehyde, 10ml absolute ethanol, 675μl sulphuric acid) and isopropanol] was added, and the solution incubated at 58°C for 30 minutes. The reaction was stopped by placing on ice for 10 minutes. The absorbance at 558 nm was measured using 200μl aliquot of the final reaction mixture in a Spectra Max M2 spectrophotometer (Molecular Devices, San Jose, CA). The absorbance values were converted into μg units using the 4-parameter standard curve generated using the standards and expressed as μg hydroxyproline/g of tissue.

### 4.10 Picrosirius red staining

Formalin fixed liver tissue was embedded in paraffin and 4μm sections were taken using a microtome. Picrosirius red staining was conducted using the standard protocol employed by the Imaging Core facility at the Oklahoma Medical Research Foundation. Briefly, formalin fixed sections were deparaffinized and stained with Picrosirius Red for one hour. Excess of Picrosirius Red was removed by rinsing in acidified water. Sections were dehydrated with ethanol and cleared with xylene. The images were taken using a Nikon Ti Eclipse microscope (Nikon, Melville, NY) for three random fields per sample and quantified using Image J software.

### 4.11 Statistical analyses

A one-way ANOVA with Tukey’s *post hoc* test was used to analyze data. All data were analyzed, and graphs were compiled using GraphPad Prism (La Jolla, CA, USA).

## ACKNOWLEDGEMENTS

The authors would like to thank Stephenson Cancer Center Tissue Pathology Core for performing H & E staining and the Imaging Core facility at the Oklahoma Medical Research Foundation for performing Picrosirius red staining. The efforts of authors were supported by NIH grants R01AG059718 (SSD), R01AG057424 (AR), and R01 AG064951 (BFM) as well as a GeroOncology Pilot Grant, (SSD), Presbyterian Health Foundation Seed grant (SSD) from the University of Oklahoma Health Sciences Center.

MML and JB were supported by NIA Training Grant T32AG052363, and MML was supported by an American Physiological Society Postdoctoral Fellowship. In addition, AR and HVR were support by the following grants from the Department of Veterans Affairs: Senior Career Research Awards 1IK6BX005238 (AR) and 1IK6 BX005234 (HVR) and Merit grants I01BX004538 (AR) and I01 BX004453 (HVR).

## CONFLICT OF INTREST

The authors declare no competing financial interests

## AUTHORS CONTRIBUTIONS

N.T. performed experiments, analyzed data, and prepared figures, R.S., S.M. and A.L.T performed western blots, E.H.N. performed the real-time qPCR, M.K and W.L. performed histopathological analysis and scoring of H&E staining sections, collaborated with B.F.M. for the hydroxyproline assay, H.V.R was responsible for designing the D+Q study and J.B., M.M.L, and A.K.B. for treating the mice with vehicle and D+Q, and A.R. and S.S.D designed the experiments and wrote and edited the manuscript as a team.

## DATA AVAILABILITY STATEMENT

The data that supports the findings of this study are available in the manuscript and supplementary material of this article. Correspondence and requests for information should be addressed to A.R or S.S.D.

