## Supplemental Table for "Senolytic Treatment Reduces Cell Senescence and Necroptosis in Sod1 Knockout Mice that is Associated with Reduced Inflammation and Hepatocellular Carcinoma"

**Supplementary Table 1S.** Changes in RT² Profiler™ PCR Arrays of 84 genes related to cytokine/chemokine for normal liver tissue from WT and Sod1KO mice treated with vehicle or D+Q.

| Gene | WT-Veh | | WT- D+Q | | Sod1KO- Veh | | Sod1KO – D+Q | | Significance difference among the groups | | | | | |
|---|---|---|---|---|---|---|---|---|---|---|---|---|---|---|
| | Fold change | SEM | Fold change | SEM | Fold change | SEM | Fold change | SEM | WT-Veh vs WT-D+Q | WT-Veh vs Sod1KO-Veh | WT-Veh vs Sod1KO-D+Q | WT-D+Q vs Sod1KO-Veh | WT-D+Q vs Sod1KO-D+Q | Sod1KO-Veh vs Sod1KO-D+Q |
| Adipoq | 1 | 0.10 | 0.26 | 0.14 | 0.08 | 0.04 | 0.09 | 0.01 | Yes | Yes | Yes | No | No | No |
| Bmp2 | 1 | 0.13 | 0.83 | 0.07 | 0.98 | 0.07 | 1.04 | 0.24 | No | No | No | No | No | No |
| Bmp4 | 1 | 0.16 | 1.01 | 0.08 | 0.62 | 0.22 | 0.71 | 0.11 | No | Yes | No | Yes | No | No |
| Bmp6 | 1 | 0.09 | 0.62 | 0.08 | 0.70 | 0.22 | 0.56 | 0.05 | Yes | Yes | Yes | No | No | No |
| Bmp7 | 1 | 0.14 | 1.15 | 0.14 | 1.02 | 0.24 | 0.53 | 0.10 | No | No | Yes | No | Yes | Yes |
| Ccl1 | 1 | 0.03 | 0.73 | 0.18 | 2.92 | 0.23 | 0.75 | 0.45 | No | Yes | No | Yes | No | Yes |
| Ccl11 | 1 | 0.16 | 0.24 | 0.13 | 0.24 | 0.07 | 0.31 | 0.06 | Yes | Yes | Yes | No | No | No |
| Ccl12 | 1 | 0.07 | 1.03 | 0.75 | 2.61 | 0.15 | 3.27 | 0.27 | No | Yes | Yes | Yes | Yes | No |
| Ccl17 | 1 | 0.24 | 0.55 | 0.26 | 1.83 | 0.72 | 0.77 | 0.10 | No | No | No | Yes | No | Yes |
| Ccl19 | 1 | 0.07 | 0.79 | 0.08 | 1.85 | 0.29 | 0.50 | 0.05 | No | Yes | Yes | Yes | No | Yes |
| Ccl2 | 1 | 0.08 | 0.83 | 0.15 | 7.63 | 1.02 | 3.02 | 0.08 | No | Yes | Yes | Yes | Yes | Yes |
| Ccl20 | 1 | 0.07 | 2.61 | 0.24 | 143.86 | 24.15 | 19.40 | 1.78 | No | Yes | No | Yes | No | Yes |
| Ccl22 | 1 | 0.14 | 0.61 | 0.27 | 1.29 | 0.14 | 1.62 | 0.20 | No | No | Yes | Yes | Yes | No |
| Ccl24 | 1 | 0.29 | 0.40 | 0.10 | 1.99 | 0.26 | 0.49 | 0.22 | Yes | Yes | Yes | Yes | No | Yes |
| Ccl3 | 1 | 0.03 | 0.40 | 0.12 | 1.03 | 0.24 | 0.78 | 0.16 | Yes | No | No | Yes | Yes | No |
| Ccl4 | 1 | 0.09 | 0.34 | 0.08 | 0.72 | 0.12 | 0.76 | 0.15 | Yes | Yes | Yes | Yes | Yes | No |
| Ccl5 | 1 | 0.07 | 1.15 | 0.38 | 1.28 | 0.22 | 2.47 | 0.23 | No | No | Yes | No | Yes | Yes |
| Ccl7 | 1 | 0.05 | 0.67 | 0.21 | 1.21 | 0.17 | 2.24 | 0.44 | No | No | Yes | No | Yes | Yes |
| Cd40lg | 1 | 0.08 | 1.05 | 0.18 | 0.63 | 0.08 | 0.55 | 0.16 | No | Yes | Yes | Yes | Yes | No |
| Cd70 | 1 | 0.25 | 0.81 | 0.13 | 0.14 | 0.06 | 0.68 | 0.38 | Yes | No | Yes | No | Yes | No |
| Cntf | 1 | 0.15 | 0.66 | 0.06 | 0.62 | 0.10 | 0.55 | 0.06 | Yes | Yes | Yes | No | No | No |
| Csf1 | 1 | 0.06 | 1.22 | 0.21 | 1.32 | 0.17 | 1.28 | 0.20 | No | No | No | No | No | No |
| Csf2 | 1 | 0.23 | 1.43 | 0.28 | 0.60 | 0.27 | 0.15 | 0.07 | No | No | Yes | Yes | Yes | No |
| Csf3 | 1 | 0.12 | 0.37 | 0.10 | 0.06 | 0.02 | 0.23 | 0.06 | Yes | Yes | Yes | Yes | No | No |
| Ctf1 | 1 | 0.19 | 0.95 | 0.12 | 0.51 | 0.12 | 0.33 | 0.09 | No | Yes | Yes | Yes | Yes | No |
| Cx3cl1 | 1 | 0.07 | 0.61 | 0.10 | 1.29 | 0.67 | 4.36 | 0.33 | No | No | Yes | No | Yes | Yes |

| | | | | | | | | | | | | | | |
|---|---|---|---|---|---|---|---|---|---|---|---|---|---|---|
| Cxcl1 | 1 | 0.15 | 0.54 | 0.07 | 25.76 | 3.99 | 8.80 | 0.84 | No | Yes | Yes | Yes | Yes | Yes |
| Cxcl10 | 1 | 0.06 | 0.91 | 0.13 | 2.25 | 0.32 | 19.41 | 1.26 | No | No | Yes | No | Yes | Yes |
| Cxcl11 | 1 | 0.12 | 0.83 | 0.20 | 0.60 | 0.15 | 0.16 | 0.01 | No | Yes | Yes | No | Yes | Yes |
| Cxcl12 | 1 | 0.03 | 0.85 | 0.07 | 1.02 | 0.24 | 0.48 | 0.08 | No | No | Yes | No | Yes | Yes |
| Cxcl13 | 1 | 0.09 | 0.88 | 0.17 | 5.08 | 0.24 | 1.40 | 0.22 | No | Yes | Yes | Yes | Yes | Yes |
| Cxcl16 | 1 | 0.05 | 0.69 | 0.06 | 1.82 | 0.32 | 2.06 | 0.30 | No | Yes | Yes | Yes | Yes | No |
| Cxcl3 | 1 | 0.11 | 0.40 | 0.17 | 1.63 | 0.19 | 2.42 | 0.24 | Yes | Yes | Yes | Yes | Yes | Yes |
| Cxcl5 | 1 | 0.12 | 2.49 | 0.10 | 4.75 | 0.62 | 1.13 | 0.16 | Yes | Yes | No | Yes | Yes | Yes |
| Cxcl9 | 1 | 0.09 | 0.39 | 0.02 | 0.76 | 0.15 | 0.39 | 0.06 | Yes | Yes | Yes | Yes | No | Yes |
| Fasl | 1 | 0.16 | 0.96 | 0.10 | 0.92 | 0.09 | 0.72 | 0.06 | No | No | Yes | No | Yes | No |
| Gpi1 | 1 | 0.09 | 0.81 | 0.09 | 3.73 | 0.10 | 1.85 | 0.30 | No | Yes | Yes | Yes | Yes | Yes |
| Hc | 1 | 0.36 | 0.75 | 0.11 | 0.88 | 0.24 | 0.62 | 0.05 | No | No | No | No | No | No |
| Ifna2 | 1 | 1.05 | 0.24 | 0.06 | 0.62 | 0.19 | 0.43 | 0.10 | No | No | No | No | No | No |
| Ifng | 1 | 0.14 | 0.29 | 0.04 | 0.41 | 0.14 | 0.68 | 0.29 | Yes | Yes | No | No | Yes | No |
| Il10 | 1 | 0.13 | 1.09 | 0.23 | 1.34 | 0.06 | 1.30 | 0.12 | No | Yes | No | No | No | No |
| Il11 | 1 | 0.06 | 0.78 | 0.21 | 0.75 | 0.31 | 0.80 | 0.11 | No | No | No | No | No | No |
| Il12a | 1 | 0.11 | 0.54 | 0.07 | 3.14 | 0.37 | 0.95 | 0.44 | No | Yes | No | Yes | No | Yes |
| Il12b | 1 | 0.10 | 1.17 | 0.05 | 0.89 | 0.20 | 2.02 | 0.30 | No | No | Yes | No | Yes | Yes |
| Il13 | 1 | 0.23 | 1.05 | 0.01 | 0.51 | 0.14 | 0.60 | 0.19 | No | Yes | Yes | Yes | Yes | No |
| Il15 | 1 | 0.21 | 1.10 | 0.21 | 1.44 | 0.31 | 1.19 | 0.18 | No | No | No | No | No | No |
| Il16 | 1 | 0.17 | 1.04 | 0.30 | 1.31 | 0.18 | 2.98 | 0.42 | No | No | Yes | No | Yes | Yes |
| Il17a | 1 | 0.20 | 0.44 | 0.06 | 0.08 | 0.02 | 0.37 | 0.13 | Yes | Yes | Yes | Yes | No | Yes |
| Il17f | 1 | 0.12 | 0.85 | 0.18 | 0.69 | 0.11 | 0.64 | 0.12 | No | Yes | Yes | No | No | No |
| Il18 | 1 | 0.10 | 0.76 | 0.05 | 0.86 | 0.22 | 0.68 | 0.11 | No | No | Yes | No | No | No |
| Il1a | 1 | 0.06 | 0.68 | 0.14 | 1.06 | 0.23 | 0.83 | 0.22 | No | No | No | Yes | No | No |
| Il1b | 1 | 0.06 | 0.54 | 0.05 | 2.37 | 0.25 | 1.33 | 0.11 | Yes | Yes | Yes | Yes | Yes | Yes |
| Il1rn | 1 | 0.05 | 0.42 | 0.07 | 2.59 | 0.08 | 4.16 | 0.33 | Yes | Yes | Yes | Yes | Yes | Yes |
| Il2 | 1 | 0.16 | 0.23 | 0.13 | 0.29 | 0.23 | 0.22 | 0.09 | Yes | Yes | Yes | No | No | No |
| Il21 | 1 | 0.10 | 0.61 | 0.21 | 0.75 | 0.15 | 1.25 | 0.17 | Yes | Yes | Yes | Yes | Yes | Yes |
| Il22 | 1 | 0.03 | 0.56 | 0.22 | 0.06 | 0.05 | 0.06 | 0.03 | Yes | Yes | Yes | Yes | Yes | No |
| Il23a | 1 | 0.06 | 0.96 | 0.27 | 0.88 | 0.30 | 2.33 | 0.42 | No | No | Yes | No | Yes | Yes |
| Il24 | 1 | 0.03 | 0.70 | 0.28 | 0.45 | 0.18 | 0.35 | 0.11 | No | Yes | Yes | No | No | No |
| Il27 | 1 | 0.17 | 1.01 | 0.27 | 1.68 | 0.28 | 1.78 | 0.57 | No | No | Yes | No | Yes | No |
| Il3 | 1 | 0.16 | 0.24 | 0.12 | 0.35 | 0.20 | 0.85 | 0.15 | Yes | Yes | No | No | Yes | Yes |

| | | | | | | | | | | | | | | |
|---|---|---|---|---|---|---|---|---|---|---|---|---|---|---|
| Il4 | 1 | 0.03 | 0.41 | 0.14 | 0.32 | 0.04 | 0.30 | 0.09 | Yes | Yes | Yes | No | No | No |
| Il5 | 1 | 0.12 | 0.45 | 0.15 | 1.75 | 0.24 | 0.63 | 0.35 | Yes | Yes | No | Yes | No | Yes |
| Il6 | 1 | 0.05 | 0.42 | 0.08 | 0.06 | 0.02 | 0.30 | 0.05 | Yes | Yes | Yes | Yes | Yes | Yes |
| Il7 | 1 | 0.06 | 0.94 | 0.05 | 1.51 | 0.43 | 1.60 | 0.43 | No | No | No | No | Yes | No |
| Il9 | 1 | 0.14 | 0.72 | 0.44 | 0.87 | 0.09 | 0.30 | 0.12 | No | No | Yes | No | No | Yes |
| Lif | 1 | 0.21 | 0.98 | 0.29 | 0.71 | 0.50 | 10.07 | 1.83 | No | No | Yes | No | Yes | Yes |
| Lta | 1 | 0.17 | 1.49 | 0.11 | 0.63 | 0.42 | 0.47 | 0.19 | No | No | Yes | Yes | Yes | No |
| Ltb | 1 | 0.07 | 2.17 | 0.30 | 1.57 | 0.16 | 2.12 | 0.27 | Yes | Yes | Yes | Yes | No | Yes |
| Mif | 1 | 0.04 | 1.33 | 0.30 | 1.64 | 0.22 | 2.44 | 0.22 | Yes | Yes | Yes | No | Yes | Yes |
| Mstn | 1 | 0.22 | 0.35 | 0.26 | 0.28 | 0.11 | 0.88 | 0.13 | Yes | Yes | No | No | Yes | Yes |
| Nodal | 1 | 0.12 | 1.10 | 0.15 | 1.02 | 0.22 | 1.61 | 0.24 | No | No | Yes | No | Yes | Yes |
| Osm | 1 | 0.18 | 0.38 | 0.05 | 0.94 | 0.45 | 2.00 | 0.32 | Yes | No | Yes | No | Yes | Yes |
| Pf4 | 1 | 0.06 | 0.89 | 0.12 | 3.95 | 0.40 | 3.31 | 0.34 | No | Yes | Yes | Yes | Yes | Yes |
| Ppbp | 1 | 0.22 | 2.45 | 0.15 | 4.50 | 1.46 | 4.03 | 0.79 | No | Yes | Yes | Yes | No | No |
| Spp1 | 1 | 0.26 | 1.20 | 0.06 | 0.90 | 0.05 | 1.48 | 0.48 | Yes | Yes | Yes | Yes | Yes | Yes |
| Tgfb2 | 1 | 0.13 | 0.88 | 0.07 | 0.99 | 0.12 | 2.49 | 0.21 | No | No | Yes | No | Yes | Yes |
| Thpo | 1 | 0.11 | 1.02 | 0.07 | 1.43 | 0.27 | 1.30 | 0.10 | No | Yes | No | Yes | No | No |
| Tnf | 1 | 0.08 | 1.30 | 0.12 | 7.86 | 0.33 | 8.90 | 0.94 | No | Yes | Yes | Yes | Yes | No |
| Tnfrsf11b | 1 | 0.05 | 1.18 | 0.13 | 4.28 | 0.17 | 1.25 | 0.23 | No | Yes | No | Yes | No | Yes |
| Tnfsf10 | 1 | 0.10 | 0.86 | 0.04 | 0.93 | 0.15 | 0.85 | 0.17 | No | No | No | No | No | No |
| Tnfsf11 | 1 | 0.53 | 0.53 | 0.18 | 0.28 | 0.03 | 0.34 | 0.12 | No | Yes | Yes | No | No | No |
| Tnfsf13b | 1 | 0.04 | 1.52 | 0.12 | 2.69 | 0.28 | 1.03 | 0.18 | Yes | Yes | No | Yes | Yes | Yes |
| Vegfa | 1 | 0.10 | 0.89 | 0.07 | 0.85 | 0.17 | 0.94 | 0.35 | No | No | No | No | No | No |
| Xcl1 | 1 | 0.04 | 1.08 | 0.16 | 1.14 | 0.10 | 1.07 | 0.10 | No | No | No | No | No | No |

**Supplementary Table 2S** - Changes in RT² Profiler™ PCR Arrays of 84 genes related to liver cancer for normal liver tissue from WT and Sod1KO mice treated with vehicle or D+Q as well as tumor tissue from Sod1KO mice treated with vehicle.

| Gene | WT-Veh | | WT-D+Q | | Sod1KO-Veh | | Sod1KO – D+Q | | Sod1KO-Tumor | | Significance difference among the groups | | | | | | |
|---|---|---|---|---|---|---|---|---|---|---|---|---|---|---|---|---|---|
| | Fold change | SEM | Fold change | SEM | Fold change | SEM | Fold change | SEM | Fold change | SEM | WT-Veh vs WT-D+Q | WT-Veh vs Sod1KO-Veh | WT vs Sod1KO-D+Q | WT vs Sod1KO-Tumor | Sod1KO-veh vs Sod1KO-D+Q | Sod1KO-veh vs Sod1KO-Tumor | Sod1KO-D+Q vs Sod1KO-Tumor |
| Adam17 | 1 | 0.09 | 0.67 | 0.05 | 0.63 | 0.06 | 0.77 | 0.11 | 0.69 | 0.08 | No | Yes | No | No | No | No | No |
| Akt1 | 1 | 0.15 | 0.78 | 0.06 | 0.69 | 0.06 | 0.64 | 0.10 | 0.88 | 0.11 | No | Yes | Yes | No | No | No | No |
| Angpt2 | 1 | 0.10 | 0.72 | 0.05 | 0.65 | 0.08 | 0.89 | 0.14 | 1.87 | 0.11 | Yes | Yes | No | Yes | Yes | Yes | Yes |
| Bax | 1 | 0.07 | 0.87 | 0.08 | 1.41 | 0.09 | 1.62 | 0.17 | 1.67 | 0.10 | No | Yes | Yes | Yes | No | Yes | No |
| Bcl2 | 1 | 0.08 | 1.01 | 0.16 | 0.59 | 0.10 | 0.90 | 0.18 | 1.00 | 0.16 | No | Yes | No | No | Yes | Yes | No |
| Bcl2l1 | 1 | 0.17 | 0.84 | 0.06 | 0.94 | 0.06 | 0.93 | 0.16 | 0.99 | 0.14 | No | No | No | No | No | No | No |
| Bid | 1 | 0.14 | 0.95 | 0.08 | 0.98 | 0.09 | 1.20 | 0.09 | 1.01 | 0.18 | No | No | No | No | No | No | No |
| Birc2 | 1 | 0.09 | 0.88 | 0.09 | 0.73 | 0.07 | 0.84 | 0.10 | 0.62 | 0.02 | No | Yes | No | Yes | No | No | Yes |
| Birc5 | 1 | 0.13 | 1.24 | 0.12 | 0.81 | 0.07 | 0.82 | 0.16 | 2.03 | 0.20 | No | No | No | Yes | No | Yes | Yes |
| Casp8 | 1 | 0.18 | 1.11 | 0.12 | 1.21 | 0.09 | 1.02 | 0.10 | 1.38 | 0.12 | No | No | No | Yes | No | No | Yes |
| Ccl5 | 1 | 0.03 | 1.08 | 0.12 | 1.05 | 0.12 | 0.96 | 0.09 | 2.00 | 0.14 | No | No | No | Yes | No | Yes | Yes |
| Ccnd1 | 1 | 0.05 | 0.95 | 0.17 | 0.35 | 0.04 | 0.85 | 0.15 | 0.59 | 0.21 | No | Yes | No | Yes | Yes | No | No |
| Ccnd2 | 1 | 0.05 | 0.96 | 0.15 | 1.21 | 0.03 | 1.08 | 0.15 | 1.34 | 0.24 | No | No | No | Yes | No | No | No |
| Cdh1 | 1 | 0.07 | 0.82 | 0.07 | 2.78 | 0.21 | 3.50 | 0.17 | 25.43 | 1.42 | No | Yes | Yes | Yes | No | Yes | Yes |
| Cdh13 | 1 | 0.07 | 0.62 | 0.03 | 0.38 | 0.04 | 0.56 | 0.10 | 0.34 | 0.11 | Yes | Yes | Yes | Yes | Yes | No | Yes |
| Cdkn1a | 1 | 0.11 | 0.52 | 0.10 | 0.59 | 0.15 | 0.89 | 0.18 | 0.87 | 0.12 | Yes | Yes | No | No | Yes | No | No |
| Cdkn1b | 1 | 0.11 | 0.78 | 0.07 | 0.61 | 0.10 | 0.71 | 0.11 | 0.47 | 0.07 | Yes | Yes | Yes | Yes | No | No | Yes |
| Cdkn2a | 1 | 0.14 | 1.14 | 0.09 | 4.15 | 0.46 | 0.98 | 0.19 | 4.57 | 0.24 | No | Yes | No | Yes | Yes | No | Yes |
| Cflar | 1 | 0.09 | 0.71 | 0.06 | 0.49 | 0.07 | 0.64 | 0.11 | 0.46 | 0.06 | Yes | Yes | Yes | Yes | No | No | Yes |
| Ctnnb1 | 1 | 0.06 | 1.04 | 0.10 | 0.86 | 0.03 | 0.90 | 0.13 | 0.78 | 0.07 | No | No | No | Yes | No | No | No |
| Cxcr4 | 1 | 0.09 | 1.36 | 0.15 | 1.42 | 0.21 | 1.53 | 0.17 | 6.26 | 0.70 | No | No | No | Yes | No | Yes | Yes |
| Dab2ip | 1 | 0.13 | 0.88 | 0.06 | 1.03 | 0.11 | 0.98 | 0.15 | 1.17 | 0.19 | No | No | No | No | No | No | No |
| Dlc1 | 1 | 0.10 | 1.14 | 0.17 | 0.66 | 0.11 | 0.73 | 0.13 | 0.46 | 0.13 | No | Yes | No | Yes | No | No | No |

| | | | | | | | | | | | | | | | | | |
|---|---|---|---|---|---|---|---|---|---|---|---|---|---|---|---|---|---|
| E2f1 | 1 | 0.11 | 0.63 | 0.11 | 0.67 | 0.08 | 0.76 | 0.19 | 0.97 | 0.21 | Yes | Yes | No | No | No | No | No |
| Egf | 1 | 0.11 | 0.98 | 0.08 | 1.06 | 0.25 | 1.04 | 0.13 | 0.51 | 0.10 | No | No | No | Yes | No | Yes | Yes |
| Egfr | 1 | 0.15 | 1.22 | 0.09 | 0.77 | 0.19 | 0.55 | 0.11 | 1.26 | 0.13 | No | No | Yes | No | No | Yes | Yes |
| Ep300 | 1 | 0.08 | 1.08 | 0.07 | 1.01 | 0.02 | 1.01 | 0.09 | 1.33 | 0.06 | No | No | No | Yes | No | Yes | Yes |
| Fadd | 1 | 0.11 | 0.88 | 0.02 | 0.88 | 0.08 | 0.84 | 0.17 | 0.74 | 0.09 | No | No | No | Yes | No | No | No |
| Fas | 1 | 0.14 | 0.76 | 0.06 | 0.69 | 0.14 | 0.87 | 0.05 | 0.48 | 0.09 | Yes | Yes | No | Yes | No | No | Yes |
| Fhit | 1 | 0.11 | 0.96 | 0.10 | 0.67 | 0.05 | 0.68 | 0.08 | 0.53 | 0.17 | No | Yes | Yes | Yes | No | No | No |
| Flt1 | 1 | 0.09 | 0.71 | 0.01 | 0.85 | 0.10 | 0.80 | 0.06 | 0.77 | 0.10 | Yes | No | Yes | Yes | No | No | No |
| Fzd7 | 1 | 0.14 | 1.09 | 0.06 | 0.60 | 0.15 | 0.89 | 0.20 | 0.53 | 0.09 | Yes | Yes | Yes | Yes | Yes | Yes | Yes |
| Gadd45b | 1 | 0.12 | 0.97 | 0.04 | 0.41 | 0.11 | 0.54 | 0.10 | 0.51 | 0.10 | No | Yes | Yes | Yes | No | No | No |
| Gstp1 | 1 | 0.18 | 1.24 | 0.14 | 2.46 | 0.25 | 2.85 | 0.77 | 4.17 | 0.70 | No | Yes | Yes | Yes | No | Yes | Yes |
| Hgf | 1 | 0.14 | 0.57 | 0.08 | 0.59 | 0.14 | 0.72 | 0.11 | 0.32 | 0.05 | Yes | Yes | Yes | Yes | No | Yes | Yes |
| Hhip | 1 | 0.06 | 0.74 | 0.07 | 0.44 | 0.10 | 0.45 | 0.13 | 0.18 | 0.12 | Yes | Yes | Yes | Yes | No | Yes | Yes |
| Hras1 | 1 | 0.06 | 0.69 | 0.06 | 0.66 | 0.14 | 0.87 | 0.12 | 0.61 | 0.07 | Yes | Yes | No | Yes | Yes | No | Yes |
| Igf2 | 1 | 0.03 | 0.60 | 0.18 | 0.46 | 0.08 | 0.60 | 0.16 | 20.61 | 3.69 | No | No | No | Yes | No | Yes | Yes |
| Igfbp1 | 1 | 0.15 | 2.00 | 0.14 | 1.51 | 0.29 | 2.22 | 0.20 | 1.56 | 0.18 | Yes | Yes | Yes | Yes | Yes | No | Yes |
| Igfbp3 | 1 | 0.06 | 0.77 | 0.10 | 0.39 | 0.07 | 0.42 | 0.18 | 0.41 | 0.20 | No | Yes | Yes | Yes | No | No | No |
| Irs1 | 1 | 0.03 | 0.98 | 0.06 | 0.68 | 0.19 | 1.21 | 0.19 | 0.45 | 0.10 | No | Yes | No | Yes | Yes | No | Yes |
| Itgb1 | 1 | 0.03 | 0.76 | 0.07 | 0.77 | 0.04 | 0.75 | 0.10 | 1.13 | 0.11 | Yes | Yes | Yes | No | No | Yes | Yes |
| Kdr | 1 | 0.06 | 0.85 | 0.03 | 0.80 | 0.12 | 0.75 | 0.13 | 0.85 | 0.07 | Yes | Yes | Yes | No | No | No | No |
| Lef1 | 1 | 0.16 | 0.43 | 0.07 | 0.91 | 0.21 | 0.91 | 0.22 | 0.54 | 0.12 | Yes | No | No | Yes | No | Yes | Yes |
| Mcl1 | 1 | 0.15 | 0.85 | 0.10 | 0.89 | 0.05 | 0.88 | 0.07 | 1.01 | 0.04 | No | No | No | No | No | No | No |
| Met | 1 | 0.12 | 1.05 | 0.16 | 1.26 | 0.13 | 0.98 | 0.13 | 1.44 | 0.24 | No | No | No | Yes | No | No | Yes |
| Msh2 | 1 | 0.12 | 0.86 | 0.08 | 0.77 | 0.11 | 0.78 | 0.15 | 0.95 | 0.14 | No | No | No | No | No | No | No |
| Msh3 | 1 | 0.05 | 0.79 | 0.08 | 0.61 | 0.10 | 0.70 | 0.18 | 0.39 | 0.04 | No | Yes | Yes | Yes | No | No | Yes |
| Mtdh | 1 | 0.11 | 0.82 | 0.08 | 0.86 | 0.06 | 0.81 | 0.11 | 1.38 | 0.18 | No | No | No | Yes | No | Yes | Yes |
| Myc | 1 | 0.08 | 1.22 | 0.17 | 2.68 | 0.21 | 1.43 | 0.03 | 6.94 | 0.86 | No | Yes | No | Yes | Yes | Yes | Yes |
| Nfkb1 | 1 | 0.13 | 0.94 | 0.04 | 0.88 | 0.04 | 0.84 | 0.07 | 1.06 | 0.07 | No | No | No | No | No | Yes | Yes |
| Nras | 1 | 0.05 | 0.76 | 0.07 | 0.86 | 0.02 | 0.76 | 0.09 | 1.07 | 0.09 | Yes | No | Yes | No | No | Yes | Yes |

| | | | | | | | | | | | | | | | | | |
|---|---|---|---|---|---|---|---|---|---|---|---|---|---|---|---|---|---|
| Opcml | 1 | 0.14 | 0.21 | 0.10 | 0.10 | 0.03 | 0.21 | 0.07 | 0.12 | 0.04 | Yes | Yes | Yes | Yes | No | No | No |
| Pdgfra | 1 | 0.09 | 0.90 | 0.13 | 1.31 | 0.30 | 1.04 | 0.13 | 0.62 | 0.09 | No | No | No | Yes | No | Yes | Yes |
| Pin1 | 1 | 0.10 | 0.88 | 0.09 | 0.78 | 0.10 | 0.89 | 0.13 | 0.64 | 0.09 | No | No | No | Yes | No | No | Yes |
| Pten | 1 | 0.10 | 0.92 | 0.07 | 0.83 | 0.07 | 0.73 | 0.10 | 0.83 | 0.08 | No | No | Yes | No | No | No | No |
| Ptgs2 | 1 | 0.11 | 0.23 | 0.15 | 0.41 | 0.18 | 0.34 | 0.13 | 1.57 | 0.37 | Yes | Yes | Yes | Yes | No | Yes | Yes |
| Ptk2 | 1 | 0.12 | 0.90 | 0.09 | 0.76 | 0.16 | 0.98 | 0.10 | 0.96 | 0.17 | No | No | No | No | No | No | No |
| Pycard | 1 | 0.04 | 1.34 | 0.11 | 1.86 | 0.37 | 1.78 | 0.25 | 1.97 | 0.22 | No | Yes | Yes | Yes | No | No | No |
| Rac1 | 1 | 0.04 | 0.77 | 0.09 | 0.72 | 0.07 | 0.73 | 0.15 | 0.77 | 0.15 | No | Yes | Yes | No | No | No | No |
| Rassf1 | 1 | 0.10 | 0.73 | 0.05 | 0.74 | 0.05 | 0.84 | 0.13 | 1.00 | 0.11 | Yes | Yes | No | No | No | Yes | No |
| Rb1 | 1 | 0.14 | 0.95 | 0.09 | 0.77 | 0.12 | 0.88 | 0.10 | 0.83 | 0.09 | No | No | No | No | No | No | No |
| Reln | 1 | 0.09 | 0.99 | 0.10 | 1.06 | 0.22 | 0.85 | 0.06 | 0.47 | 0.10 | No | No | No | Yes | No | Yes | Yes |
| Rhoa | 1 | 0.13 | 0.89 | 0.08 | 0.69 | 0.02 | 0.79 | 0.10 | 0.69 | 0.02 | No | Yes | Yes | Yes | No | No | No |
| Runx3 | 1 | 0.08 | 0.90 | 0.03 | 0.70 | 0.19 | 0.90 | 0.05 | 0.97 | 0.11 | No | Yes | No | No | No | Yes | No |
| Sfrp2 | 1 | 0.12 | 0.26 | 0.09 | 0.10 | 0.05 | 0.25 | 0.07 | 0.06 | 0.02 | Yes | Yes | Yes | Yes | No | No | Yes |
| Smad4 | 1 | 0.16 | 0.73 | 0.04 | 0.75 | 0.09 | 0.66 | 0.08 | 0.94 | 0.04 | Yes | Yes | Yes | No | No | No | Yes |
| Smad7 | 1 | 0.18 | 2.13 | 0.16 | 1.29 | 0.18 | 1.49 | 0.31 | 1.17 | 0.06 | Yes | No | Yes | No | No | No | No |
| Socs1 | 1 | 0.11 | 0.97 | 0.18 | 0.66 | 0.17 | 1.09 | 0.14 | 1.42 | 0.29 | No | No | No | Yes | Yes | Yes | No |
| Socs3 | 1 | 0.15 | 0.90 | 0.15 | 4.43 | 0.76 | 1.70 | 0.33 | 5.02 | 0.39 | No | Yes | No | Yes | Yes | No | Yes |
| Stat3 | 1 | 0.08 | 1.09 | 0.19 | 1.54 | 0.19 | 1.34 | 0.12 | 1.22 | 0.18 | No | Yes | No | No | No | No | No |
| Tcf4 | 1 | 0.14 | 0.67 | 0.11 | 0.57 | 0.11 | 0.78 | 0.15 | 0.80 | 0.17 | Yes | Yes | No | No | No | No | No |
| Tert | 1 | 0.17 | 0.95 | 0.08 | 0.75 | 0.17 | 1.13 | 0.25 | 0.45 | 0.02 | No | No | No | Yes | Yes | No | Yes |
| Tgfa | 1 | 0.05 | 0.72 | 0.05 | 1.18 | 0.13 | 1.29 | 0.18 | 1.88 | 0.31 | No | No | No | Yes | No | Yes | Yes |
| Tgfb1 | 1 | 0.17 | 0.74 | 0.06 | 1.01 | 0.13 | 0.95 | 0.18 | 1.49 | 0.20 | No | No | No | Yes | No | Yes | Yes |
| Tgfbr2 | 1 | 0.03 | 1.54 | 0.13 | 3.07 | 0.20 | 1.82 | 0.14 | 6.65 | 0.89 | No | Yes | No | Yes | Yes | Yes | Yes |
| Tlr4 | 1 | 0.14 | 0.89 | 0.12 | 1.05 | 0.17 | 0.95 | 0.24 | 1.67 | 0.21 | No | No | No | Yes | No | Yes | Yes |
| Tnfrsf10b | 1 | 0.10 | 0.74 | 0.09 | 9.23 | 0.53 | 9.16 | 0.80 | 23.7 | 3.01 | Yes | No | No | No | Yes | No | No |
| Tnfsf10 | 1 | 0.16 | 0.75 | 0.02 | 0.66 | 0.08 | 0.85 | 0.11 | 0.72 | 0.10 | Yes | Yes | No | Yes | No | No | No |
| Trp53 | 1 | 0.16 | 0.82 | 0.06 | 0.85 | 0.16 | 1.03 | 0.11 | 0.96 | 0.11 | No | No | No | No | No | No | No |
| Vegfa | 1 | 0.11 | 0.91 | 0.08 | 0.60 | 0.13 | 0.97 | 0.25 | 0.56 | 0.13 | No | Yes | No | Yes | Yes | No | Yes |

| | | | | | | | | | | | | | | | | | |
|---|---|---|---|---|---|---|---|---|---|---|---|---|---|---|---|---|---|
| Wt1 | 1 | 0.08 | 0.66 | 0.13 | 4.19 | 0.26 | 1.21 | 0.17 | 0.50 | 0.16 | No | Yes | No | Yes | Yes | Yes | Yes |
| Xiap | 1 | 0.08 | 0.87 | 0.08 | 0.72 | 0.07 | 0.68 | 0.11 | 0.79 | 0.07 | No | Yes | Yes | Yes | No | No | No |
| Yap1 | 1 | 0.11 | 0.69 | 0.05 | 0.75 | 0.05 | 0.76 | 0.09 | 0.60 | 0.08 | Yes | Yes | Yes | Yes | No | No | No |

**Supplementary Table 3S.** List of Real Time - PCR primers used in the study

| Gene | Forward Sequence | Reverse Sequence |
|---|---|---|
| Cdkn2a(p16$^{Ink4a}$) | 5’-CCCAACGCCCCGAACT-3’ | 5’-GCAGAAGAGCTGCTACGTGAA-3’ |
| Cdkn1a(p21$^{Cip1}$) | 5’-GTCAGGCTGGTCTGCCTCCG-3’ | 5’-CGGTCCCGTGGACAGTGAGCAG-3’ |
| p53 | 5’-GTATTTCACCCTCAAGATCC-3’ | 5’-TGGGCATCCTTTAACTCTA-3’ |
| IL-6 | 5’-TGGTACTCCAGAAGACCAGAGG-3’ | 5’-AACGATGATGCACTTGCAGA-3’ |
| IL-1α | 5’-AGGGAGTCAACTCATTGGCG-3’ | 5’-TGGCAGAACTGTAGTCTTCGT-3’ |
| IL-1β | 5’-AGGTCAAAGGTTTGGAAGCA-3’ | 5’-TGAAGCAGCTATGGCAACTG-3’ |
| CXCL-1 | 5’-ACCCGCTCGCTTCTCTGT-3’ | 5’-AAGGGAGCTTCAGGGTCAAG-3’ |
| CXCL-2 | 5’-CCTGGTTCAGAAAATCATCCA-3’ | 5’-CTTCCGTTGAGGGACAGC-3’ |
| CXCL-8 | 5’-AGACAGCAGAGCACACAAGC-3’ | 5’-ATGGTTCCTTCCGGTGGT-3’ |
| CXCL-10 | 5’-CCAAGTGCTGCCGTCATTTTC-3’ | 5’-GGCTCGCAGGGATGATTTCAA |
| GDF-15 | 5′-GTTAGCCAAAGACTGCCACTG-3′ | 5′-CCTTGAGCCCATTCCACA-3′ |
| PAI-1 | 5’-GACACCCTCAGCATGTTCATC-3’ | 5’-AGGGTTGCACTAAACATGTCAG-3’ |
| MMP3 | 5’GTTGGAGAACATGGAGACTTTGT-3’ | 5’-CAAGTTCATGAGCAGCAACCA-3’ |
| MMP12 | 5’-TGCACTCTGCTGAAAGGAGTCT-3’ | 5’-GTCATTGGAATTCTGTCCTTTCCA-3’ |
| H2A.X | 5’-ACGAGGAGCTCAACAAGCTG-3’ | 5’-GTGGCGCTG GTCTTCTTG-3’ |
| F4/80 | 5'-CCCCAGTGTCCTTACAGAGTG-3' | 5'-GTGCCCAGAGTGGATGTCT-3' |
| HPRT | 5’-CTGGTGAAAAGGACCTCTCG-3’ | 5’-TGAAGTACTCATTATAGTCAAGGGCA-3’ |
| β-microglobulin | 5′-CACTGACCGGCCTGTATGC-3′ | 5′-GGGTGGCGTGAGTATACTTGAAT-3′ |
| β-actin | 5′-ATGGATGACGATATCGCTG-3′ | 5′-GTTGGTAACAATGCCATGTTC-3′ |

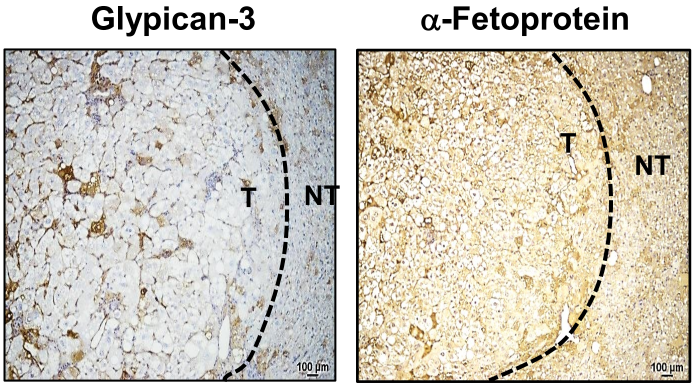


Figure 1S Immunohistochemistry (IHC) analysis of glypican-3 and α-fetoprotein in liver tissue from Sod1KO treated with vehicle. Tumor regions (T) are demarked from non-tumor (NT) regions with dotted line. Scale bar: 100 µm.
